# Host-parasite coevolution in populations of constant and variable size

**DOI:** 10.1101/012435

**Authors:** Yixian Song, Chaitanya S. Gokhale, Andrei Papkou, Hinrich Schulenburg, Arne Traulsen

## Abstract

The matching-allele and gene-for-gene models are widely used in mathematical approaches that study the dynamics of host-parasite interactions. Agrawal and Lively (Evolutionary Ecology Research 4:79-90, 2002) captured these two models in a single framework and numerically explored the associated time discrete dynamics of allele frequencies. Here, we present a detailed analytical investigation of this unifying framework in continuous time and provide a generalization. We extend the model to take into account changing population sizes, which result from the antagonistic nature of the interaction and follow the Lotka-Volterra equations. Under this extension, the population dynamics become most complex as the model moves away from pure matching-allele and becomes more gene-for-gene-like. While the population densities oscillate with a single oscillation frequency in the pure matching-allele model, a second oscillation frequency arises under gene-for-gene-like conditions. These observations hold for general interaction parameters and allow to infer generic patterns of the dynamics. Our results suggest that experimentally inferred dynamical patterns of host-parasite coevolution should typically be much more complex than the popular illustrations of Red Queen dynamics. A single parasite that infects more than one host can substantially alter the cyclic dynamics.

## 1 Introduction

The antagonistic interaction between hosts and their parasites are of particular interest in ecology and evolution because they are ubiquitous and usually associated with high selection pressure that affects numerous life history traits. Because of the negative effect of parasites on host fitness, the study of these interactions is of central importance in biomedical (Woolhouse et al., 2002, 2005), agricultural (Van der Plank, 1984; Gladieux et al., 2011) and species conservation research (Altizer et al., 2003; Thompson et al., 2010). The exact dynamics are usually evaluated with the help of mathematical models. Among these, the models including an explicit genetic description of host-parasite interaction, such as gene-for-gene (GfG) and matching-allele (MA) models, are particularly widespread. Genetic interaction is usually incorporated by taking into account the current understanding of resistance-infectivity patterns in biological systems. The gene-for-gene (GfG) model was proposed by Flor (1956) to capture disease resistance patterns in plants. Here, a host individual carrying a resistance gene can recognize parasites harboring the corresponding avirulence product and trigger a defense response averting the infection (Jones and Dangl, 2006). Inspired by self-nonself recognition in immune systems (Grosberg and Hart, 2000), the matching-allele (MA) model was introduced to reflect host-pathogen interactions in animals. In this case, parasites carrying a certain allele can only invade host individuals with the corresponding allele. By combining predictive power of mathematical modeling and their connection to the empirical data, these models successfully served to understand key evolutionary problems. To mention only the most important examples, these models were used to assess the Red Queen hypothesis for the evolution of sexual reproduction (Lively, 2010), the maintenance of genetic diversity by parasite-mediated selection (Lively and Apanius, 1995), and the role of the cost of resistance/virulence in coevolution (Leonard, 1977; Parker, 1994).

Agrawal and Lively (2002) developed a general model that interpolates between a pure matching-allele model and a pure gene-for-gene model, as a single parameter is tuned between 0 and 1. This model was introduced for haplotypes of two loci with mutation and recombination. Variance in host and parasite allele frequency was plotted as an evaluation of the time discrete dynamics. The highly dynamical aspects of matching-allele models were observed across most of the MA-GfG continuum. Agrawal and Lively showed that cyclic dynamics of host and parasite genotypes is observed not only in the MA model, but also in all the intermediate models and in the GfG model. This finding indicates that the Red Queen theory for the evolution of sex does not hinge upon the use of a particular model for host parasite interactions. However, this study was computational and only performed for particular parameter sets due to the complexity of the model. Instead of tackling the dynamics from an analytical perspective to allow for general statements for all parameter sets, subsequent theoretical approaches have increased complexity of the assumed interaction in order to increase the biological realism, for instance by defining a multi-locus model that deals with various combinations of MA loci and GfG loci (Agrawal and Lively, 2003).

The aim of our study is to improve our understanding of host-parasite coevolution by focusing on an analytical characterization of the involved dynamics. We investigate both the impact of different types of interaction and the consequence of interaction-dependent population size changes. We simplify the model of Agrawal and Lively and focus on a single locus to keep interaction among loci from interfering with the conclusion, in particular the differences between the GfG model (Tellier and Brown, 2007a) and the MA model (Sardanyés and Solé, 2008). We use the assumptions of Agrawal and Lively (2002) inspired by Parker (1994) to connect the two popular models by a single parameter, but also provide an alternative, linear interpolation in the discussion. To enhance clarity, we focus on a system with two host and two parasite genotypes and use their interaction to characterize the involved evolutionary dynamics. In addition, we depart from the usual assumption of constant population size and apply the Lotka-Volterra equations to acknowledge inter-dependent population dynamics during host-parasite coevolution. To compare the dynamics with a model assuming constant population density, we apply the Replicator Dynamics with the same interaction matrix between hosts and parasites. While the dynamics between the two models is different, it seems to be crucial to understand both constant as well as changing population size, as there are biological examples for both of them.

We conducted a linear stability analysis at the interior fixed point of the resulting nonlinear dynamical system, which indicates critical differences in dynamical patterns between the models of host-parasite coevolution. Either with constant or with changing population density, the population densities oscillate with a single frequency in a pure MA model. In a model deviating from MA, a second oscillation frequency arises with changing population density, but not with constant population size.

## 2 Model

We consider haploid hosts and parasites with two alleles on a single locus. Hence, there are two host types and two parasite types that are denoted by 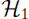, 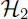, 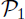, and 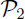, respectively. In the simplest case, each parasite type can only infect the corresponding host type. Hence, no host/parasite type is superior to the other. This case corresponds to the matching-allele model, which under the assumption of constant population density is equivalent to the evolutionary game of matching pennies (Hofbauer and Sigmund, 1998; Traulsen et al., 2005).

In a GfG model, the virulent parasite 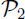 can potentially infect both hosts, the one with susceptible allele 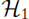 and the one with resistance allele 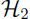. Yet, the avirulent parasite 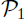 can only infect the susceptible host 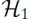, as the host 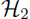 with the resistance allele can prevent infection by 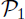. Thus, there is an advantage to the virulent parasite and the resistant host. To maintain the different types in the population, intrinsic costs of virulence and resistance have been suggested (Leonard, 1994).

Fig. 1 illustrates the fitness of the two parasites on each host for the MA and the GfG model and also for two intermediate cases, where the parasite 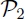 can “partially” infect the host 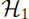.

**Figure 1:**
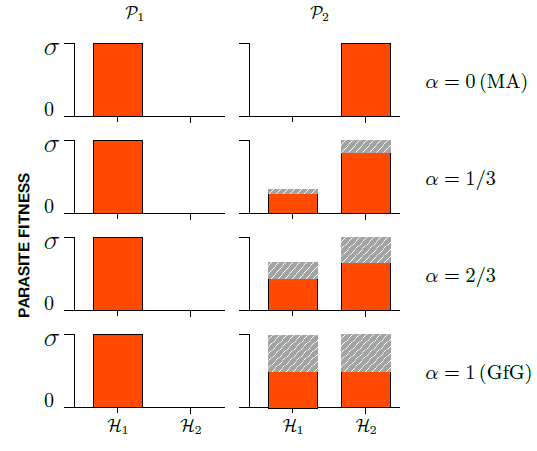
Fitness of avirulent parasite 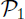 and virulent parasite 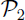 on the two hosts 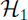 and 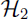 for the matching-allele model (*α* = 0, top), the gene-for-gene model (*α* = 1, bottom), and two intermediate models (*α* = 1/3 and *α* = 2/3). Gray areas represent the fitness reduction for 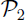 due to the cost of virulence *κ* = 1/2, which is *ακσ* in 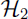 (Eq. (1b)), hence, *σ*/2 in GfG model. In 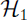 the fitness reduction for 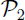 due to the cost of virulence is *α*^2^*κσ*.

We simplified the model of Agrawal and Lively (2002) by regarding only one locus. The interactions between hosts and parasites can be expressed with two matrices (corresponding to a bi-matrix game in evolutionary game theory). For the parasite, we assume that the interactions with the hosts increase birth rates. The fitness effects arising from the interactions of the parasite with the host are given by the matrix

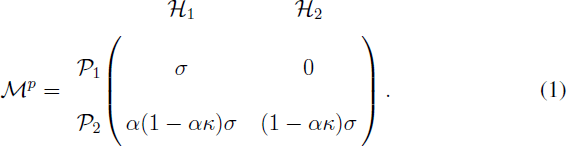

The maximum virulence of the parasite is given by *σ*. The parameter *κ* describes the cost for the parasite virulence, as usually assumed in the GfG model. This model interpolates between the MA and the GfG model as the parameter *α* is varied between 0 and 1.

For the host, we assume that these interactions increase the death rate according to the matrix

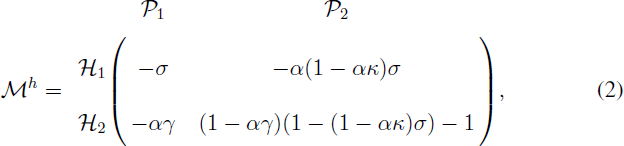

where the parameter *γ* describes the cost for the host resistance.

We assume a large population size and focus on the change in population densities. The population densities of the two host and two parasite types are given by *h*_1_, *h*_2_, *p*_1_, and *p*_2_, respectively. The population dynamics of the hosts and parasites can be captured by a set of differential equations,

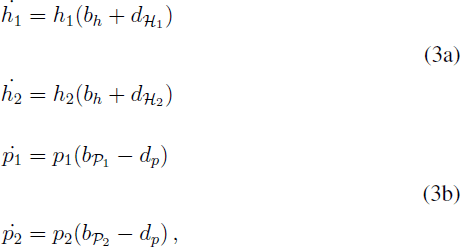

where *b*_*h*_ is the birth rate of both hosts, and *d*_*p*_ is the death rate of both parasites. As discussed above, the death rates of the hosts and the birth rates of the parasites are directly affected by host-parasite interactions. From the interaction matrices Eqs. (1) and (2), the death rates for the hosts and the birth rates for the parasites are given by

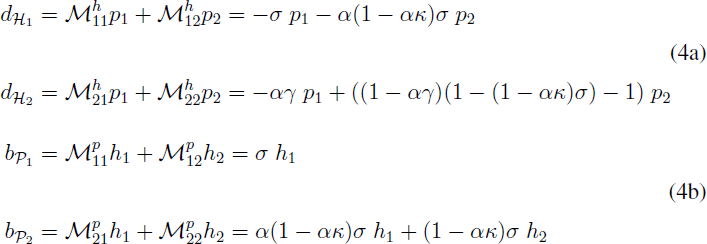

We will choose the host birth rate *b*_*h*_ and parasite death rate *d*_*p*_ in two distinct ways. Our first approach assumes constant values for *b*_*h*_ and *d*_*p*_, which leads to a host/parasite population that is changing in size. This corresponds to the standard Lotka-Volterra dynamics. The second approach focuses on relative abundances of host and parasite alleles and implies a normalization of the population size. This corresponds to the Replicator Dynamics in evolutionary game theory, which implies constant population size in our context.

### Changing population size induced by interactions

With constant host birth rate *b*_*h*_ and parasite death rate *d*_*p*_, inserting the host parasite interactions Eqs. (4) into the dynamical system Eqs. (3) leads to

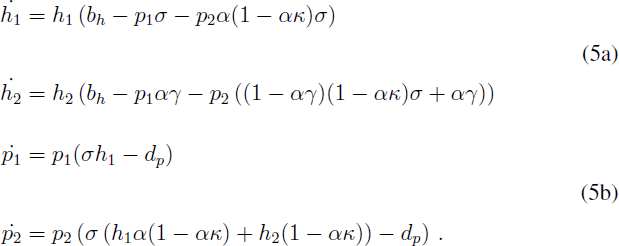

This model results in changes in the population sizes of both hosts and parasites. In particular, the changes are caused by the antagonistic interactions between the hosts and the parasite - as a consequence of the Lotka-Volterra relationship.

### Constant population size

To obtain a model of constant population size that is comparable to the one described above, we retain the interaction matrices and adjust the host birth rate and parasite death rate to maintain the population size. Requiring constant *h*_1_ + *h*_2_ and constant *p*_1_ + *p*_2_ implies 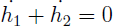 as well as 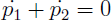. This leads to

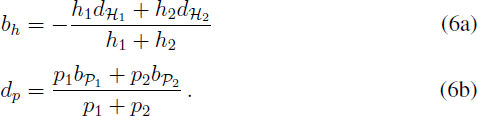

The normalization *h*_1_ + *h*_2_ = 1 implies that a single equation for *h*_1_ is sufficient to describe the dynamics for the host. Similarly, due to the normalization *p*_1_ + *p*_2_ = 1 the parasite dynamics are fully captured by tracking *p*_1_. Applying the dynamical host birth and parasite death rates in the dynamical system Eqs. (3), the equations become identical to the Replicator Dynamics (RD) (Hofbauer and Sigmund, 1998; Taylor and Jonker, 1978; Schuster and Sigmund, 1983),

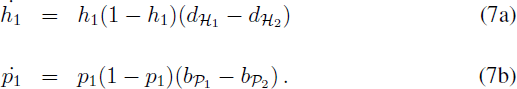

While the death rates of the host still depend on the parasites and the birth rates of the parasites still depend on the hosts, the dynamics of this system is in general less complex than in the case of changing population size, as it is only two-dimensional.

## 3 Population dynamics

To obtain first information about the population dynamics, we calculated the trajectories of the system numerically for a particular set of parameters. In addition, we identify the fixed points of the differential equations and study their stability to gain insight into the coevolutionary dynamics for all parameter sets. More specifically, we can use a linear stability analysis of the unique interior fixed point to infer the dynamical patterns arising in this system (Strogatz, 2000; Tellier and Brown, 2007a). Finally, we also assess constants of motion.

### 3.1 Numerical solution of the dynamics

To illustrate the differences in the population dynamics described in Eqs. (5) and (7), we show numerical solutions side by side in Fig. 2.

**Figure 2:**
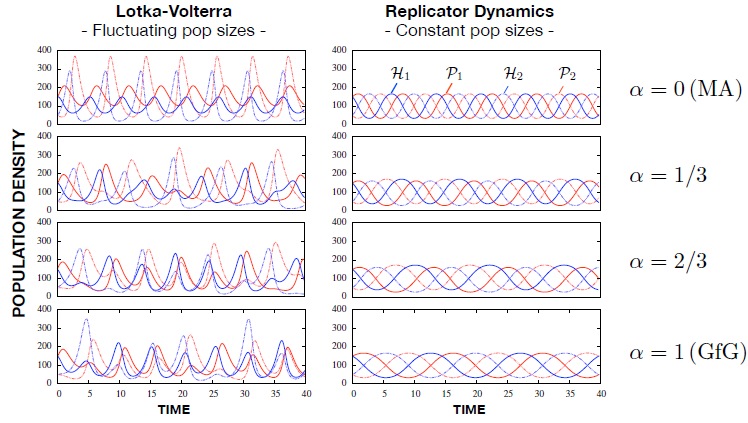
Example of population dynamics based on the Lotka-Volterra equations (left) and the Replicator Dynamics (right). While the dynamics on the right side resembles the common Red Queen pattern, the left side is more complex. In a pure matching-allele model (top), the plot on the left shows two independent sets of Lotka-Volterra dynamics, one for 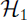 and 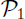 (blue and red solid lines, correspondingly) and a second one for 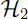 and 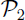 (blue and red dotted lines). As the model deviates from MA model with increasing *α* (rows 2-4) more complicated dynamics arise, since the four population densities of 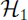, 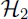, 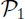, and 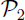 are coupled (parameters *γ* = 0.005, *κ* = 0.5, and *σ* = 0.01 for both Lotka-Volterra and Replicator Dynamics. Host birth rate *b*_*h*_ = 1.5 and parasite death rate *d*_*p*_ = 1.0 in the Lotka-Volterra case. Initial population densities *h*_1_ = *p*_1_ = 150, *h*_2_ = *p*_2_ = 50).

The dynamics in models with constant host and parasite population sizes resemble the common Red Queen pattern. Under changing population sizes the system is uncoupled into two independent host-parasite pairs in a pure MA model. As the model deviates from the MA model with increasing *α*, the dynamics becomes more complex, since the four population densities of the types 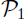, 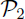, 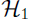, and 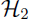 are coupled.

### 3.2 Stability of boundary fixed points

The fixed points of the system are the points where all population sizes remain constant in time, 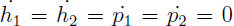. The position of the fixed points and their stability change with changing parameters.

For the Lotka-Volterra dynamics, a trivial fixed point is (*h*_1_, *h*_2_, *p*_1_, *p*_2_) = (0, 0, 0, 0) where both the hosts and parasites are absent, cf. Eqs. (5). Additionally, extinction of one host and the associated parasite leads to two further fixed points, 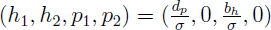 and 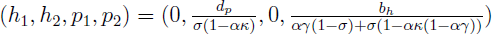. In gene-for-gene-like models, *α* > 0, the susceptible host 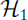 and the virulent 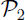 can coexist in the absence of 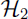 and 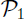, 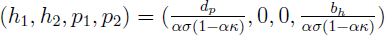. The opposite case, coexistence between 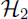 and 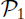 in the absence of 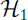 and 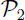 is not possible, as our host-parasite interaction model assumes that the birth rate of 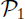 is zero in the absence of 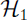. A linear stability analysis of the Lotka-Volterra model shows that all boundary fixed points are unstable for *αγ* < *σ*. That is, if the cost of resistance *αγ* (which is scaled by the amount of GfG influence) is less than the maximum host fitness reduction caused by infection *σ*, then all host and parasite types will coexist.

The Replicator Dynamic system, Eq. (7), has four fixed points at the boundaries, each is reflecting fixation of one host and one parasite: (*h*_1_, *p*_1_) = (0, 0), (*h*_1_, *p*_1_) = (1, 0), (*h*_1_, *p*_1_) = (0, 1), (*h*_1_, *p*_1_) = (1, 1). A linear stability analysis reveals that all these fixed points are unstable.

### 3.3 Stability of the interior fixed point

In addition to the boundary fixed points, the system has a unique fixed point in the interior. In the Lotka-Volterra system, we obtain a non-trivial fixed point of the four dimensional dynamical system described in Eqs. (5) when *αγ* < *σ*. This fixed point, where all types coexist, is given by

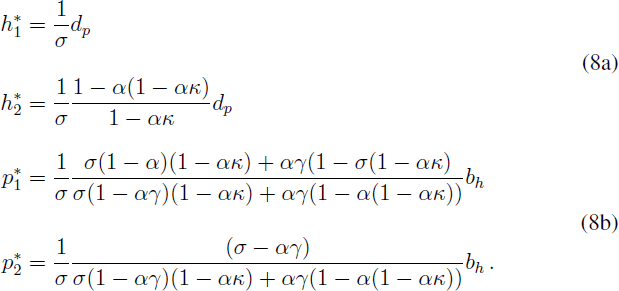

For *αγ* > *σ*, the resistant host is always disadvantageous because of the high cost of the resistance allele (*γ*). Consequently, extinction of 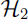 and 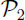 then becomes a stable fixed point. For *αγ* < *σ*, 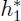 and 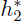 increase linearly with parasites’ death rate *d*_*p*_, while 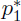 and 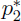 increase linearly with hosts’ birth rate *b*_*h*_. A linear stability analysis of the interior fixed point (see Appendix A for details) shows that the equilibrium is neutrally stable. Close to the interior fixed point, the system exhibits undamped oscillations. More specifically, the four eigenvalues of the Jacobi-matrix are two distinct pairs of complex conjugates without real parts. This means there are two distinct oscillation frequencies in the system,

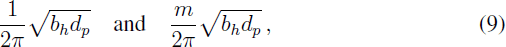

where

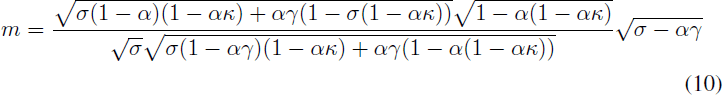

measures the ratio between the two oscillation frequencies. This ratio decreases when we move away from the MA interaction model. For *α* ≈ 0, we find

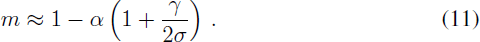

In particular, for the MA model both oscillation frequencies collapse into a single one. However, all solutions for *α* > 0 exhibit both of the frequencies (Fig. 3).

**Figure 3:**
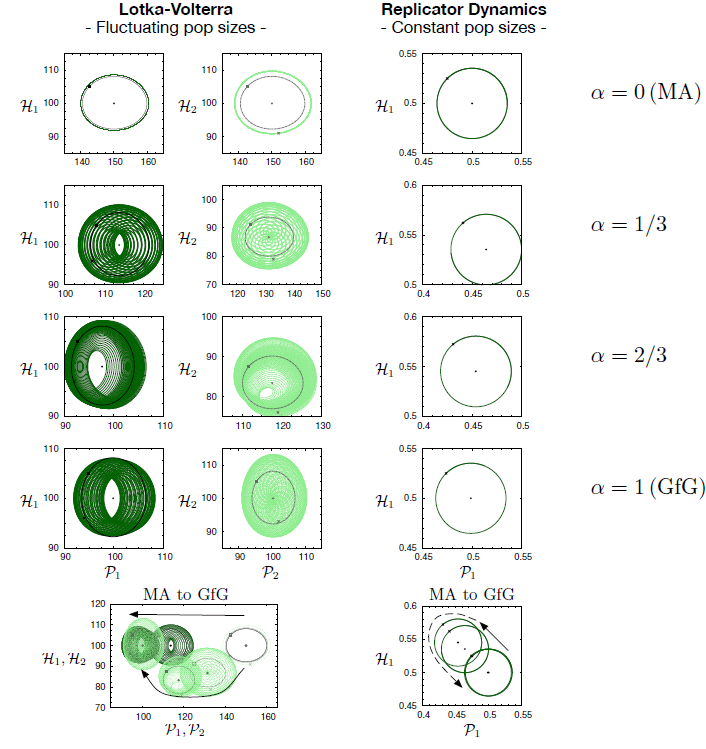
Trajectories close to the interior fixed points (black points) on the *h*_1_ – *p*_1_ plane (dark green solid lines both for LV and RD equations) and the *h*_2_ – *p*_2_ plane (light green dashed lines LV only). The black crosses mark the initial conditions. The black rectangle represent a special set of initial condition while the black solid/dashed lines show the corresponding trajectories. With Replicator Dynamics the *h*_1_ – *p*_1_ trajectory is a closed circle. With Lotka-Volterra dynamics, the trajectories are closed circle when the initial conditions fulfill Eq. (24) (black lines). For the closed circles (black in LV and green in RD) the initial host population densities, *h*_1_ and *h*_2_ are 5% above the corresponding fixed point, while the parasite population densities are 5% beneath the fixed point. Except for *α* = 0 (MA) the green trajectories with LV resemble tori instead of closed circles, an implication for two oscillation frequencies. To show the shift of the interior fixed point as *α* increases from 0 to 1, the trajectories are plotted all in the same coordinate system at the bottom.

For the Replicator Dynamics system in which the population size is constant, the non-trivial fixed point of Eqs. (7) is given by

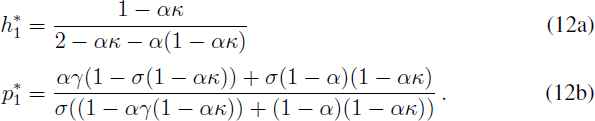

A linear stability analysis shows that the interior fixed point is again neutrally stable, as the two eigenvalues are a pair of purely imaginary, complex conjugated numbers when *αγ* < *σ* (see Appendix B for details). Hence, there is only one characteristic oscillation frequency of the dynamical system at the fixed point,

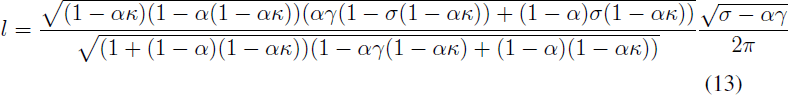

*l* has a maximum value, *σ*/(4*π*), in the pure matching-allele model (*α* = 0). Close to the matching-allele model, *α* ≈ 0, the oscillation frequency decreases with increasing *α* as

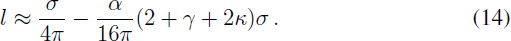

The solutions around the fixed point exhibit the oscillation frequency described by Eq. (13). The trajectories are closed circles as shown on the right side of Fig. 3.

### 3.4 Disentangling evolutionary and ecological dynamics

To clarify the ecological effect on the dynamics, particularly at the interior fixed point, we derive the dynamics of the host and parasite population sizes, *h* = *h*_1_ + *h*_2_ and *p* = *p*_1_ + *p*_2_, and the relative abundance of 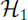 and 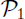 in the population, *x* = *h*1/*h* and *y* = *p*1/*p*, from Eqs. (3). According to Eqs. (3) the differential equations for the population sizes of hosts *h* and parasites *p* are

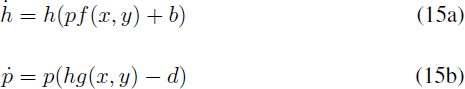

where

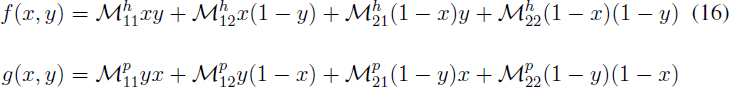

and the differential equations for relative abundances of 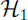 and 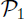 are

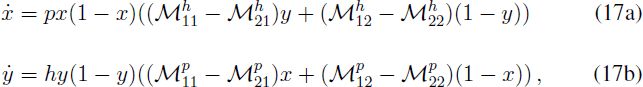

If *f*(*x*, *y*) and *g*(*x*, *y*) are constant in time Eqs. (15) yield simple Lotka-Volterra dynamics, while Eqs. (17) result in Replicator Dynamics with rescaled time if the population sizes are kept constant.

At the interior fixed point one of the oscillation frequencies, 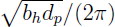, results solely from Lotka-Volterra dynamics. The other oscillation frequency

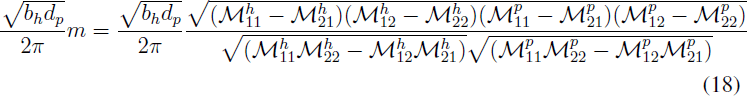

(see Eq. (35) in Appendix C) is the product of the oscillation frequency with constant population size

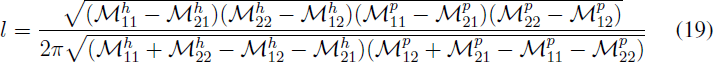

(see Eq. (39) in Appendix C) and the geometric mean of host and parasite population size 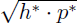, i.e.,

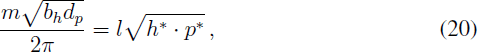

with

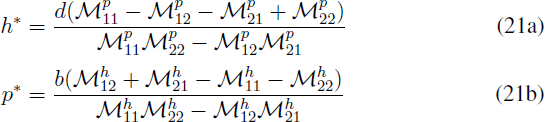

(calculated from Eqs. (32) in Appendix C). Thus, one of the oscillations results purely from ecological interactions, while the other one arises from the combination of ecology and evolution in our system.

### 3.5 Constants of motion

The system with constant population size has a constant of motion (Eq. (10.22) in (Hofbauer and Sigmund, 1998)) given by

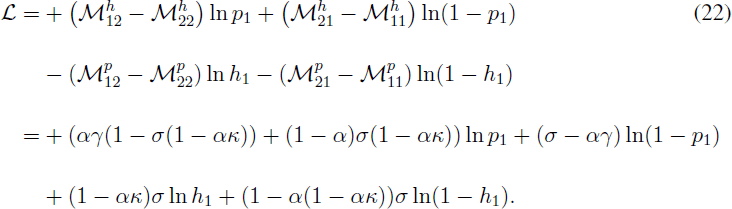

Due to 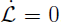, we obtain sustained oscillations for any initial condition, even far away from the interior fixed point Eq. (12)

The case of changing population size is more intricate. In the case of a matching allele model *α* = 0, the two equations decouple and we have two independent Lotka-Volterra systems with sustained oscillations, characterized by the two constants of motion

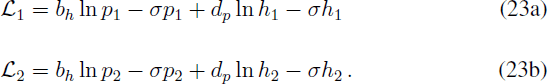

While we do not find a constant of motion for the general case of *α* > 0, particular initial conditions can lead to invariants. If the initial condition fulfills

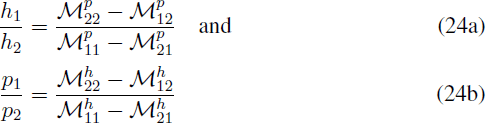

which corresponds to a two-dimensional subspace, then there are two constants that remain invariant over time,

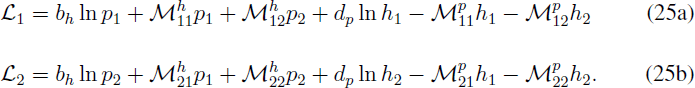

Note that with the condition Eq. (24a) the ratio *p*_1_/*p*_2_ remains constant and with the condition Eq. (24b), the ratio *h*_1_/*h*_2_ remains constant. This shows that the nature of the dynamics in this case does not only depend on the choice of parameters, but also on the initial state of the system, which in principle leads to a further complication for the corresponding experimental systems.

## 4 Discussion

### 4.1 Short overview

Host-parasite interactions are acknowledged as a driving evolutionary force promoting biological diversity and sexual reproduction (Lively and Apanius, 1995; Lively, 2010), with the MA and GfG model being the most popular models to describe the genetic interaction for coevolving hosts and parasites (Frank, 1993b; Otto and Michalakis, 1998; Lively, 2009; Gokhale et al., 2013; Luijckx et al., 2013; Clay and Kover, 1996; Brown and Tellier, 2011). Despite a number of important insights provided within their framework, the generality of findings often suffers from the complexity of the models employed and, as a consequence, the difficulty to fully understand them analytically (Bergelson et al., 2001).

In this study, we present a very general yet parsimonious model of host-parasite coevolution spanning from MA to GfG with either constant or interaction-driven changing population size. Derived analytical solutions revealed that the coevolution dynamics differs qualitatively between the models with constant and changing population sizes. Apart from the pure MA situation, the well known Red Queen dynamics with trajectories on closed circles is only observed in models with constant population size. This implies that the patterns of host-parasite dynamics to be expected in real biological systems can be much more intricate than suggested by the most popular theoretical models.

### 4.2 Main results and analytical solution

Our study is based on a simplification of the model suggested by Agrawal and Lively (2002) that explores a continuum between the MA and GfG models. We study the model in the context of haplotypes with a single locus, but relax the restriction to constant population size. With a coevolutionary system of two host and two parasite types we achieved an analytical characterization across the entire parameter space. To study ecological effects caused by the victim-exploiter interaction (Tellier and Brown, 2007b) between hosts and parasites, we consider models with changing population size aside of models with constant population size. Under the assumption of constant population size, the dynamics in MA and GfG models appear to be very similar, both showing sustained oscillations with only one oscillation frequency. Yet, introducing changing population size according to the Lotka-Volterra equations, we obtain distinct patterns of the population dynamics. For changing population sizes, a single oscillation frequency is present only in the MA model. An additional oscillation frequency arises for all other points on the MA-GfG continuum in that case. In other words, changing population size leads to a much more complex dynamics in GfG-like models, but not in the pure MA model.

In Gokhale et al. (2013) the analysis of allele fixation time for the MA model revealed that Lotka-Volterra dynamics in combination with the associated stochastic effects quickly break down the Red Queen circle. As the dynamics in GfG-like models take a completely different nature with changing population size, the influence of Lotka-Volterra dynamics on the Red Queen circle is yet unclear and remains to be assessed in more detail in the future, especially as our current analysis did not take stochastic effects into account.

### 4.3 Generality of results

To test the generality of our findings we additionally analyzed the interaction matrix suggested by Parker (1994) (Eqs. (36)). There a factor that denotes the fitness reduction of the avirulent parasite encountering the resistant host and an advantage of the virulent parasite meeting the resistant host are assumed in addition. These two parameters together with the costs of resistance and virulence determine whether the model is MA or GfG. Again we obtain two distinct oscillation frequencies for the population dynamics with changing population sizes in GfG-like models (the ratio is shown in Eq. (37) in the Appendix C).

Despite the convincing biological relevance of the interaction matrix elements in (Agrawal and Lively, 2002), they do not change monotonically on the MA-GfG continuum, e.g., with a cost of virulence *κ* > 0.5, 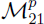 in Eq. (1) first increases then decreases as *α* increases from 0 to 1. As an alternative interpolation, we therefore also considered interaction matrices that describe a linear transition from MA to GfG model, such that

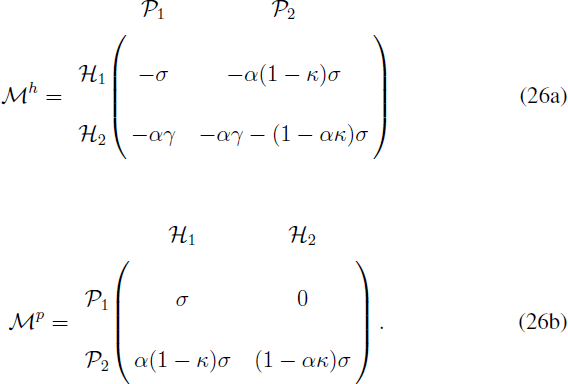

The analysis in Appendix D shows that our conclusion also holds for the linear interpolation. One should keep in mind that both MA and GfG models and even the intermediate models proposed by Parker or Agrawal & Lively or us are only a small subset of the possible models for host-parasite interaction. An observation that will hold for any such model is that as long as the population sizes are kept constant, the population dynamics follows a closed circle with a single oscillation frequency. However, with changing population size a second oscillation frequency arises when the model become GfG-like, which can lead to much more intricate dynamics. For a pure MA model or an inverse MA model (where the diagonal instead of the off-diagonal matrix elements are zero), there still is only one oscillation frequency (see Eqs. (35) in Appendix C).

### 4.4 Impact of eco-evo feedback in genetically explicit models

In the last two decades it has been realized that evolutionary changes can be faster than previously thought and, thus, occurring on the same time-scale as ecological interactions, especially in case of coevolving hosts and parasites (Hendry and Kinnison, 1999; Thompson, 1998; Hairston et al., 2005; Schoener, 2011). Population dynamics can influence the pace of coevolution via so called eco-evolutionary feedbacks, or even give rise to a new type of coevolutionary dynamics as we showed in our study. Interestingly enough, a comprehensive part of the theoretical studies on eco-evolutionary feedbacks is conducted within the framework of game theory and adaptive dynamics (Hofbauer and Sigmund, 1998; Dieckmann, 2002). In contrast to our model, these approaches usually do not include an explicit definition of genetic interaction between the species, which limits their application for interpreting patterns of genetic variability in natural populations (Day, 2005). Rapid changes in genetic composition may lead to perturbation in host demography and disease dynamics, as was observed for the myxoma virus epidemic in Australian populations of European rabbit (Fenner and Fantini, 1999). Genetic adaptation can improve overall population fitness and “buffer” the unfavorable impact of pathogens (evolutionary rescue) (Gomulkiewicz and Holt, 1995). However population perturbations may constrain adaptability, for example, via enhancing inbreeding, affecting trait heritabilities and disturbing allele composition irrespective of natural selection (O’Brien and Evermann, 1988; Lande, 1988; Gomulkiewicz and Houle, 2009; Saccheri and Hanski, 2006). Thus, models accounting simultaneously for the genetic basis of host-parasite interaction and associated population dynamics may be necessary to fully understand ongoing coevolution among species and the effect it would have on genetic diversity. We are aware of only a few such models (Frank, 1991, 1993a; Gandon et al., 1996; Quigley et al., 2012; Gokhale et al., 2013; Ashby and Gupta, 2014), and most of them confirm that ecological parameters can have a very strong effect on coevolution.

### 4.5 Implications for maintenance of genetic diversity

Numerous field studies identified the presence of comprehensive heritable variation in resistance-infectivity patterns for plant and animal populations and their respective pathogens, suggesting that coevolution acts to maintain genetic diversity (Van der Plank, 1984; Thompson and Burdon, 1992; Lively and Apanius, 1995; Carius et al., 2001; Wilfert and Jiggins, 2010; Luijckx et al., 2012). However, already the first studies, which attempted to explain such variation by cycling dynamics, encountered the problem of stability. This is especially true for the GfG model as a parasite with the virulent allele would be quickly fixed, unless having a cost of virulence (Jayakar, 1970; Leonard, 1977; Van der Plank, 1984). In addition to the cost, other factors have been examined for their potential role in maintaining variation, including epidemiological feedback (May and Anderson, 1983; Ashby and Gupta, 2014), spatial structure (Frank, 1993a; Gandon et al., 1996; Thrall and Burdon, 1997, 2002), genetic drift (Salathé et al., 2005), diffuse multi-species coevolution (Karasov et al., 2014), models with multiple alleles and multiple loci (Sasaki, 2000; Salathé et al., 2005; Tellier and Brown, 2007a). Several studies proposed that multiple factors need to act jointly for long-term coexistence of multiple resisto- and infectotypes (Bergelson et al., 2001). The view of a multifactorial basis of the maintenance of diversity creates an additional challenge for theoretical and empirical studies to disentangle them. As opposed to that, Tellier and Brown (2007b) presented a simple GfG framework showing that the general condition for stability is the presence of direct frequency-dependent selection (where fitness of an allele declines with increasing frequency of that allele itself). In this context, the distinction is made between direct frequency dependence and indirect frequency-dependent selection where fitness is mediated by the frequency of the corresponding antagonist. Direct frequency-dependent selection can be introduced in the model by incorporation of epidemiological or ecological factors (Brown and Tellier, 2011, Table 1). If we introduce a direct frequency-dependent element by applying competitive Lotka-Volterra equations or the concept of empty spaces (Hauert et al., 2006) (implying the existence of a carrying capacity) into our model, the neutrally stable interior fixed point becomes stable. Instead of forming tori or moving along closed circles, the deterministic trajectory spirals inwards. In this case, the oscillation of allele frequencies lasts longer in stochastic simulations, hence the polymorphic state is more stable.

The stability analysis derived the condition for coexistence *αγ* < *σ*, suggesting that departing from the GfG end of the continuum would increase a range of parameters at which the oscillation of allele frequencies is maintained. Therefore, patterns of “partial” infectivity by a virulent parasite are more likely to result in cycling dynamics compared to a pure GfG situation. Agrawal and Lively (2002) came to the same conclusion by evaluating computational simulations. This reinforces the importance of exploring dynamics for intermediate points on the MA-GfG continuum, especially as experimental studies provide some examples of such types of interaction (García-Arenal and Fraile, 2013). In contrast to (Tellier and Brown, 2007b) and many other studies (Agrawal and Lively, 2002; Thrall and Burdon, 2002; Tellier and Brown, 2007a), our model is implemented on a continuous time-scale and, therefore, covers host and parasite systems with overlapping generations. Interestingly, it has been proposed that models with discrete generations would favor coevolutionary cycling by synchronizing ecological and epidemiological processes (Ashby and Gupta, 2014), while in (Tellier and Brown, 2007b) the condition for stable cycling is more restrictive for discrete generations when compared to the continuous model.

## 5 Summary

In summary, we have shown that only a small and possibly biased subset of possible host-parasite interaction dynamics is captured by the mathematical models that assume fixed population size or particular genetics for the interaction, such as the MA model. Even in a simple model that allows for a full analytical description, the dynamics can vary substantially between subsequent coevolutionary cycles. We showed analytically that the complex dynamics found for changing population sizes is not a result of choosing a particular interaction matrix. The complex pattern is not limited to the set of models considered here, but rather a general property of models beyond fixed population size. Our findings highlight the importance of the interconnectedness between coevolution and population dynamics, and its potential role in understanding the generation and maintenance of genetic variation.

## Acknowledgements

YS, CSG, and AT acknowledge generous funding by the Max Planck Society. AP was funded through grants of the German Science Foundation to HS (DFG grants SCHU 1415/8 and SCHU 1415/9 within the German priority programme SPP1399 on host-parasite coevolution). AP was additionally supported by the International Max-Planck Research School (IMPRS) for Evolutionary Biology.

## A Stability of the interior fixed point in the Lotka-Volterra dynamics

In order to analyse the system at the interior fixed point 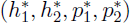, we first linearise the system around this point. For general points (*h*_1_, *h*_2_, *p*_1_, *p*_2_), the linearised system is given by by the Jacobian matrix *J*(*h*_1_, *h*_2_, *p*_1_, *p*_2_) =

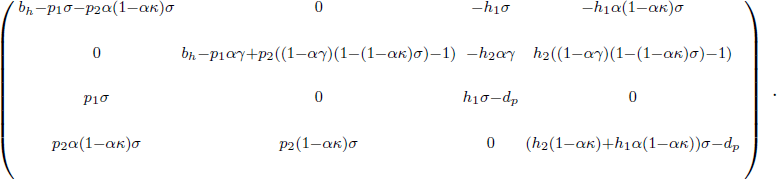

At the interior fixed point 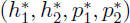, we have 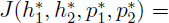

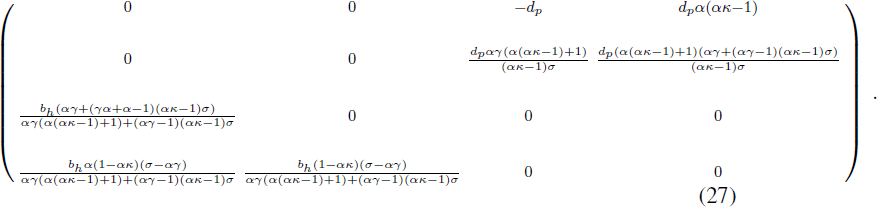

The eigenvalues of this matrix determine linear stability at the fixed point (Strogatz, 2000). If there is at least one eigenvalue with positive real part, the point would be unstable. If all eigenvalues have negative real parts, the point would be stable. In our case, the four eigenvalues are

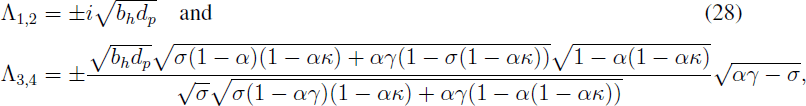

Except the term 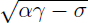, the remaining factors in in Λ_3,4_ are positive. For *αγ* > *σ*, allele 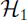 is always beneficial. Consequently, the fixed point is unstable as one of the eigenvalues Λ_3_ or Λ_4_ is positive. For *αγ* < *σ*, the fixed point is a center with neutral stability as all eigenvalues are purely imaginary. Only the case of *αγ* < *σ* is of further interest in this manuscript, as the result is straightforward in the opposite case.

## B Stability of the interior fixed point in the Replicator Dynamics

For the system with constant population size, the Jacobian matrix in general is *J*(*h*_1_, *p*_1_) =

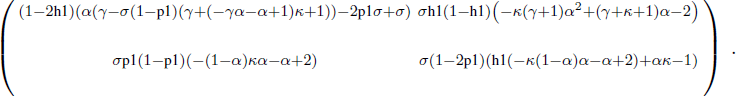

At the interior fixed point 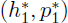, the matrix is given by 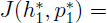

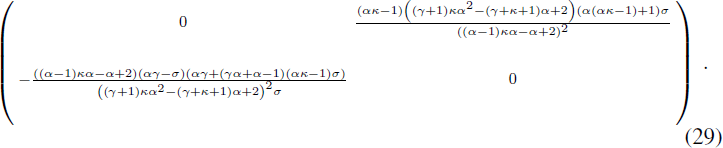

The eigenvalues are

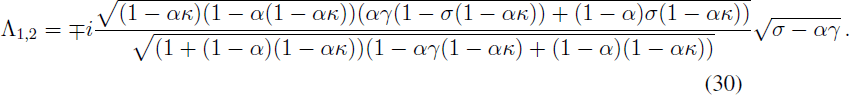

For *αγ* < *σ*, the eigenvalues are purely imaginary, hence, the fixed point is a neutral center.

## C Stability of the interior fixed point for general interaction matrices

The appearance of the second oscillation frequency at the interior fixed point in gene-for-gene-like models with changing population sizes does not depend on the exact choice of the interaction matrices in Eq. (2). To show this, we recalculate the interior fixed point and apply linear stability analysis on interaction matrices of a general form,

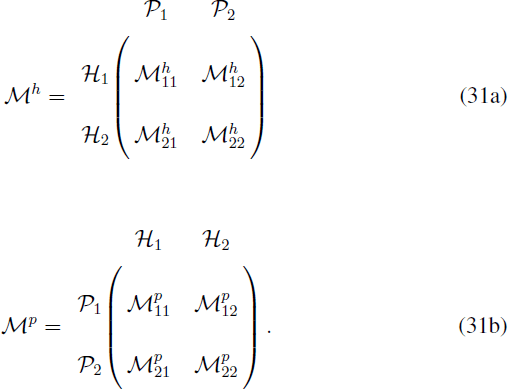

The interior fixed point for our host parasite system with Lotka-Volterra dynamics (Eq. (3)) is then

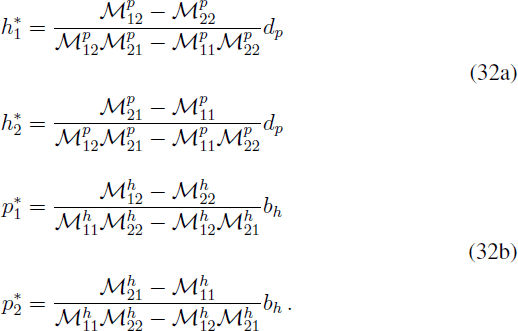

The Jacobian matrix at any defined point is *J*(*h*_1_, *h*_2_, *p*_1_, *p*_2_) =

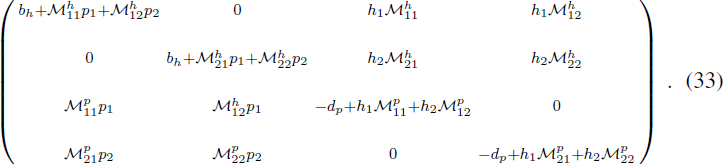

At the interior fixed point 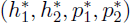, we now have

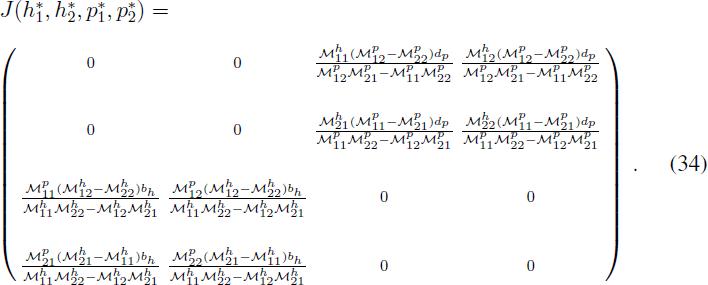

There are four eigenvalues

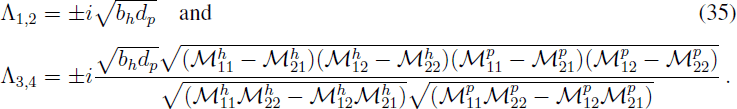

It is often assumed that (i) 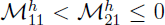 (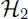 is beneficial if there is only 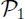 in the population), (ii) 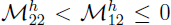 (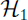 is beneficial if there is only 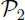 in the population), (iii) 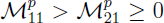 (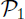 is beneficial if there is only 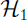 in the population), and (iv) 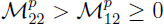 (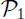 is beneficial if there is only 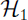 in the population). With these minimal assumptions the eigenvalues are pure imaginary, i.e., the interior fixed point is a neutrally stable center. The ratio between the eigenvalues, which determines the oscillation frequencies at the center, differs in different interaction models. For example, in Parker (Parker, 1994) the interaction matrices for haploid types are

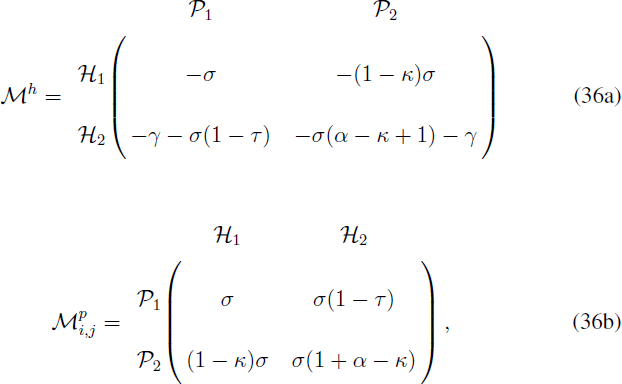

where the notations *a*, *c*, *k*, *t*, and *s* in (Parker, 1994) are changed to *α*, *γ*, *κ*, *τ*, and *σ*, respectively. According to Parker (1994), the fitness of the “narrowly virulent pathogen” 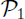 is reduced by a factor *τ* by interacting with the resistant host 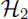; a fitness penalty *κ* (the cost of virulence) is inflicted on the “broadly virulent pathogen” 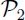 independent of which host it exploits; *α* the “advantage of adapted pathogens on resistant host” measures a special advantage of 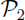 on 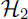; a fitness penalty *γ* (the cost of resistance) is paid by the resistant host 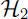. When *τ* = *κ* = *α* = 1 and *γ* = 0 the fitnesses conform to the pattern of pure MA model. When *τ* = 1 and *α* = 0 the fitnesses revert to a pure GfG pattern. The ratio between the two oscillation frequencies at the interior fixed point is

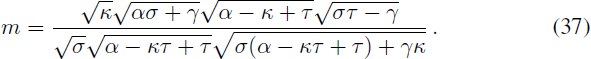

The ratio is 1 for pure MA model. With a set of parameter used in (Parker, 1994), *α* = 0.33, *γ* = 0, *κ* = 0.05, and *σ* = *τ* = 1 the ratio is about 0.1.

The same method can be applied for the system with constant population size. There the interior fixed point expressed by the general interaction matrices elements is

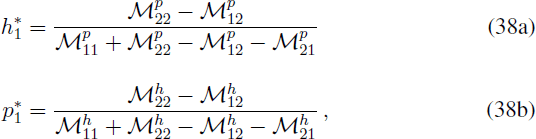

while 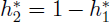 and 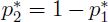. The eigenvalues of the Jacobian matrix at the interior fixed point are

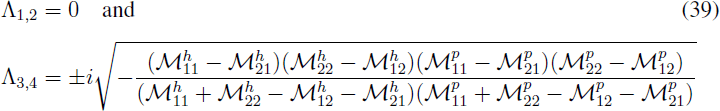

Hence, there only is one oscillation frequency at the interior fixed point in models with constant population size, regardless of the specific assumption for the interaction matrices.

## D Linear interpolation between MA and GfG models

Alternatively to the models of Agrawal and Lively (2002) and Parker (1994), one could also use a linear interpolation between MA and gene-for-gene model, where the matrix elements linearly spans over the values of the two models as a single parameter *α* varies between 0 and 1

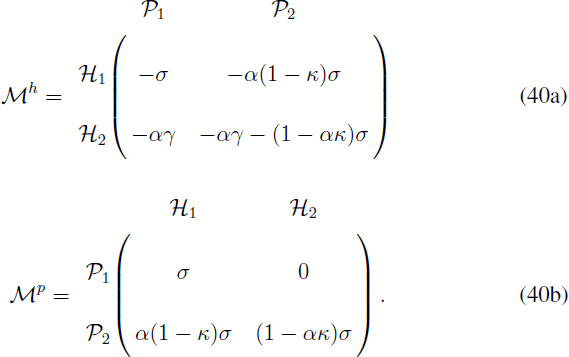

The fixed point with Lotka-Volterra dynamics is then

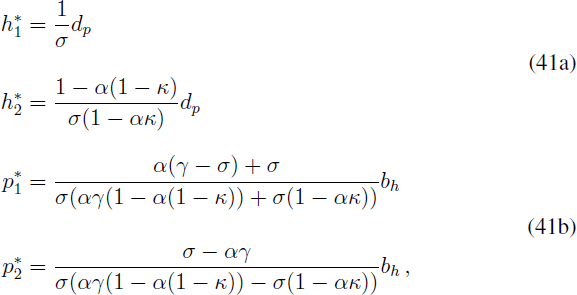

and the eigenvalues of the Jacobian matrix at this point are

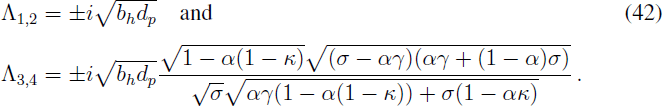

As long as *αγ* < *σ* the ratio *m* = Λ_3,4_/Λ_1,2_ increases with increasing cost of virulence *κ* while *m* decreases with increasing *α*. For *α* ≈ 0, we find

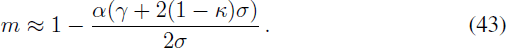

Hence, there are always two distinct oscillation frequencies at the interior fixed point in gene-for-gene-like models with changing population size.

With Replicator Dynamics the interior fixed point is

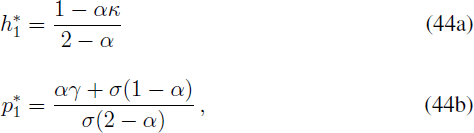

while 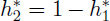 and 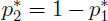. The eigenvalues of the Jacobian matrix at the interior fixed point are

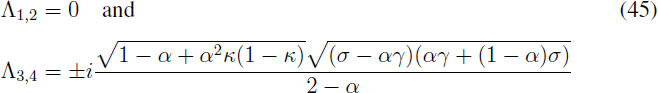

Hence, there is only one oscillation frequency *l* = Λ_3_/(*i*2*π*) at the interior fixed point in models with constant population size. As long as *αγ* < *σ*, the oscillation frequency *l* decreases with *α* and increases with *γ* and *σ*, while *l* increases with *κ* until *κ* reaches the values 1/2, then *l* decreases as *κ* increases from 1/2 to 1. For *α* ≈ 0,

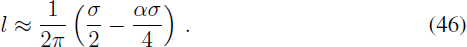

